# A non-canonical role for glutamate decarboxylase 1 in cancer cell amino acid homeostasis, independent of the GABA shunt

**DOI:** 10.1101/2021.02.17.431489

**Authors:** Bozena Samborska, Takla Griss, Eric H. Ma, Nicholas Jones, Kelsey S. Williams, Alexey Sergushichev, Radia M. Johnson, Ekaterina Esaulova, Ekaterina Loginicheva, Breanna Flynn, Julianna Blagih, Maxim N. Artyomov, Russell G. Jones, Emma E. Vincent

## Abstract

Glutamate decarboxylase 1 (GAD1) is best known for its role in producing the neurotransmitter γ-amino butyric acid (GABA) as part of the “GABA shunt” metabolic pathway, an alternative mechanism of glutamine anaplerosis for TCA cycle metabolism (Yogeeswari et al., 2005). However, understanding of the metabolic function of GAD1 in non-neuronal tissues has remained limited. Here, we show that GAD1 supports cancer cell proliferation independent of the GABA shunt. Despite its elevated expression in lung cancer tissue, GAD1 is not engaged in the GABA shunt in proliferating non-small cell lung cancer (NSCLC) cells, but rather is required for regulating amino acid homeostasis. Silencing GAD1 promotes a broad deficiency in amino acid uptake, leading to reduced glutamine-dependent TCA cycle metabolism and defects in serum- and amino acid-stimulated mTORC1 activation. Mechanistically, GAD1 regulates amino acid uptake through ATF4-dependent amino acid transporter expression including *SLC7A5* (LAT1), an amino acid transporter required for branched chain amino acid (BCAA) uptake. Overexpression of LAT1 rescues the proliferative and mTORC1 signalling defects of GAD1-deficient tumor cells. Our results, therefore, define a non-canonical role for GAD1, independent of its characterised role in GABA metabolism, whereby GAD1 regulates amino acid homeostasis to maintain tumor cell proliferation.

---

Cancer cells rely on TCA cycle metabolites to produce biosynthetic intermediates (e.g., lipids, amino acids, and nucleotides) as a means to support burgeoning proliferative rates (De Berardinis and Chandel, 2016). However, continuous depletion of TCA cycle intermediates limits mitochondrial function and proliferative capacity. To prevent this, the TCA cycle is constantly replenished by a process called anaplerosis (Owen et al., 2002). In canonical TCA cycle anaplerosis, glutamate produced from glutamine is converted to α-ketoglutarate (αKG) either by transamination reactions or by glutamate dehydrogenase (GDH/*GLUD1*) (Moreadith and Lehninger, 1984). Alternative mechanisms of glutamine anaplerosis have also been characterised. For example, expression of mitochondrial glutamate-pyruvate transaminase 2 (GPT2) has been shown to contribute to cell growth by maintaining TCA cycle anaplerosis upon inhibition of glutaminase (GLS)—the first enzyme in the canonical route for glutamine anaplerosis (Kim et al., 2019). The malate-aspartate shuttle has also been identified as an anaplerotic source of carbon and a means to adapt to loss of the TCA cycle enzyme oxoglutarate dehydrogenase (OGDH) (Allen et al., 2016).

A third metabolic pathway supporting TCA cycle anaplerosis is the GABA shunt (Figure 1A). In this pathway, glutamate is converted to GABA by GAD1, followed by transamination of GABA by GABA transaminase (ABAT) to generate succinyl-semialdehyde (SSA); SSA is finally converted to succinate by SSA-dehydrogenase (SSADH) (Yogeeswari et al., 2005). This has been proposed as an alternate route for succinate production in macrophages to induce inflammatory signalling through HIF-1α and IL-1β (Tannahill et al., 2013). Neuroendocrine cells have also been shown to use the GABA shunt in response to metabolic stress (Ippolito and Piwnica-Worms, 2014). Several studies have reported that GAD1—the rate-limiting enzyme in the GABA shunt—is overexpressed in cancers (Kimura et al., 2013; Lee et al., 2016; Maemura et al., 2003; Matuszek et al.; Yan et al., 2016) such as NSCLC (Tsuboi et al., 2019), suggesting links between GAD1 and cancer development. Through a survey of *GAD1* gene expression across several tumor types in The Cancer Genome Atlas (TCGA), we confirm elevated *GAD1* expression in several cancer types including lung adenocarcinoma (LUAD) and lung squamous cell carcinoma (LUSC) tissues (Figure 1B), with high expression of GAD1 also associated with worse survival in patients with lung cancer (logrank P = 1.6e-05) (Figure 1C). By contrast, there is little evidence of increased RNA expression of other GABA shunt enzymes—*ABAT* or *ALDH5A1* (SSADH)—in tumor tissue (Figure S1A), and increased expression of these enzymes was not associated with poor survival (Figures S1B). Previous studies have broadly hypothesized that the importance of GAD1 to tumor cells lies in its role in the GABA shunt pathway; however, this hypothesis had not been fully tested.

**Figure 1:**
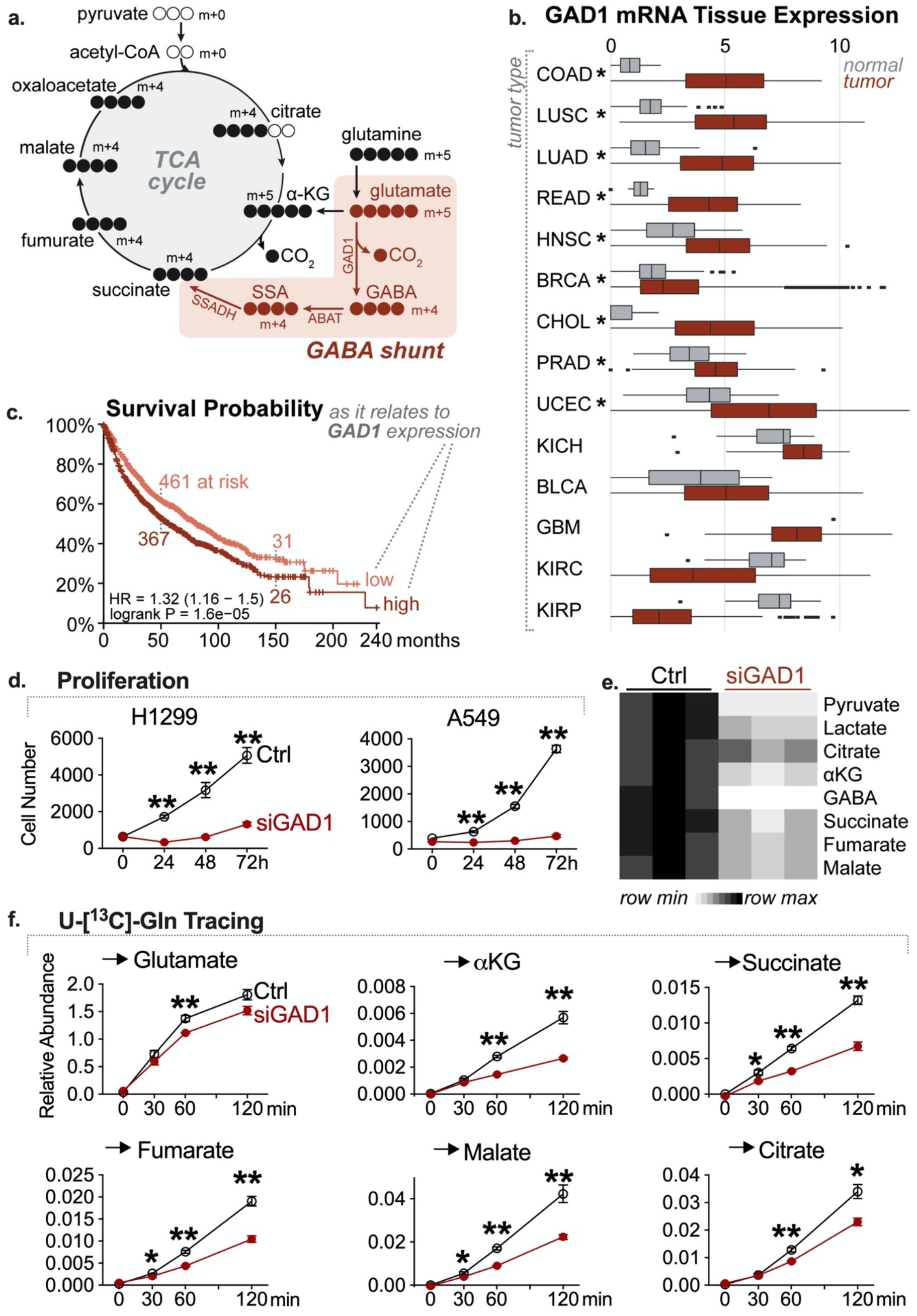
GAD1 is expressed in lung cancer and supports NSCLC proliferation and metabolism. **A**. Schematic of the GABA shunt and carbon labelling partners from U-[^13^C]-Gln. αKG: α-ketoglutarate, GAD1: glutamate dehydrogenase, ABAT: 4-aminobutyrate (GABA) aminotransferase, SSADH: succinate semi-aldehyde dehydrogenase (encoded by *ALDH5A1*). **B**. Boxplot of data taken from the The Cancer Genome Atlas (TCGA) database representing tumor tissue and normal tissue mRNA expression of GAD1. Data presented as Log2 expression. Data is sorted by t-test statistical value, with large up-regulation in tumors at the top (* denotes adjusted p < 0.01). COAD: Colon Adenocarcinoma, LUSC: Lung Squamous Cell Carcinoma, LUAD: Lung Adenocarcinoma, READ: Rectum Adenocarcinoma, HNSC: Head-Neck Squamous Cell Carcinoma, BRCA: Breast Invasive Carcinoma, CHOL: Cholangiocarcinoma, PRAD: Prostate Adenocarcinoma, UCEC: Uterine Corpus Endometrial Carcinoma, KICH: Kidney Chromophobe, BLCA: Urothelial Bladder Carcinoma, GBM: Glioblastoma, KIRC: Kidney Renal Clear Cell Carcinoma, KIRP: Cervical Kidney renal papillary cell carcinoma. **C**. Kaplan–Meier overall survival curve in lung cancer patients stratified into high (n=962) or low (n=964) expression of GAD1 mRNA using the median expression value as the cut-off point. The hazard ratio (HR) with 95% confidence intervals (CIs) and log-rank p values are displayed. **D**. Proliferation of A549 and H1299 cells transfected with either control (Ctrl) or siRNA targeting GAD1 (siGAD1). Data are represented as mean (± SD) of cell number over time for replicates (n=5). **E**. Heat map of intracellular metabolite levels in H1299 cells transfected with either control siRNA (Ctrl) or siRNA targeting GAD1 (siGAD1). Values for each metabolite abundance are normalized to a myristic acid internal standard and scaled by row. **F**. ^13^C-Glutamine utilization by control or GAD1-deficient cells. H1299 cells were transfected as in E, followed by incubation with 2 mM U-[^13^C]-Gln. Cell extracts were harvested at the time points shown, and relative abundance of ^13^C incorporation into indicated metabolites is shown. Data are represented as mean ± SEM for three independent cultures. *, p < 0.05, **, p < 0.01.

Having previously shown that NSCLC cell lines are dependent on glutamine to support cellular bioenergetics and proliferation when glucose is limiting (Vincent et al., 2015), we investigated whether these tumor cells could engage the GABA shunt as an alternative mechanism of glutamine anaplerosis. We used RNAi-mediated knockdown to silence GAD1 expression, which reduced both *GAD1* expression (Figure S1C) and GABA abundance (Figure S1D) in NSCLC cells. Silencing *GAD1* substantially inhibited the proliferation of glutamine-dependent NSCLC cell lines H1299 and A549 (Figure 1D). Metabolite levels for intermediates of both glycolysis (pyruvate, lactate) and the TCA cycle (citrate, α-ketoglutarate, succinate, fumarate, malate) were decreased in H1299 cells with GAD1 knockdown (Figure 1E and S1E), suggesting a negative impact of GAD1 depletion on central carbon metabolism. Using stable isotope labelling (SIL) with U-[^13^C]-glutamine, we found that glutamate production from ^13^C-glutamine was not persistently affected by silencing of GAD1 (Figure 1F); however, incorporation of ^13^C-glutamine-derived carbon into TCA cycle intermediates, particularly metabolites downstream of glutamate (i.e., α-ketoglutarate, succinate, and fumarate) were significantly decreased in H1299 cells lacking GAD1 (Figure 1F).

Given that GAD1 commits glutamate towards GABA biosynthesis, we hypothesized that GABA itself may function as an anaplerotic fuel for TCA cycle metabolism. Yet, despite detecting high amounts of U-[^13^C]-GABA uptake by the tumor cells tested, we observed no incorporation of U-[^13^C]-GABA-derived carbon into intermediates of the TCA cycle (Figure 2A and S2A). These data strongly suggest that NSCLC cells do not engage the GABA shunt, and that the impact of GAD1 knockdown on tumor cell proliferation may be uncoupled from its canonical enzymatic activity. Consistent with this, despite GABA having been previously shown to have pro-proliferative effects on cancer cells (Liu et al., 2008; Roberts et al., 2009; Takehara et al., 2007; Young and Bordey, 2009), exogenous GABA did not confer a proliferative advantage to A549 cells (Figure S2B) and failed to rescue cell proliferation upon GAD1 knockdown (Figure 2B).

**Figure 2:**
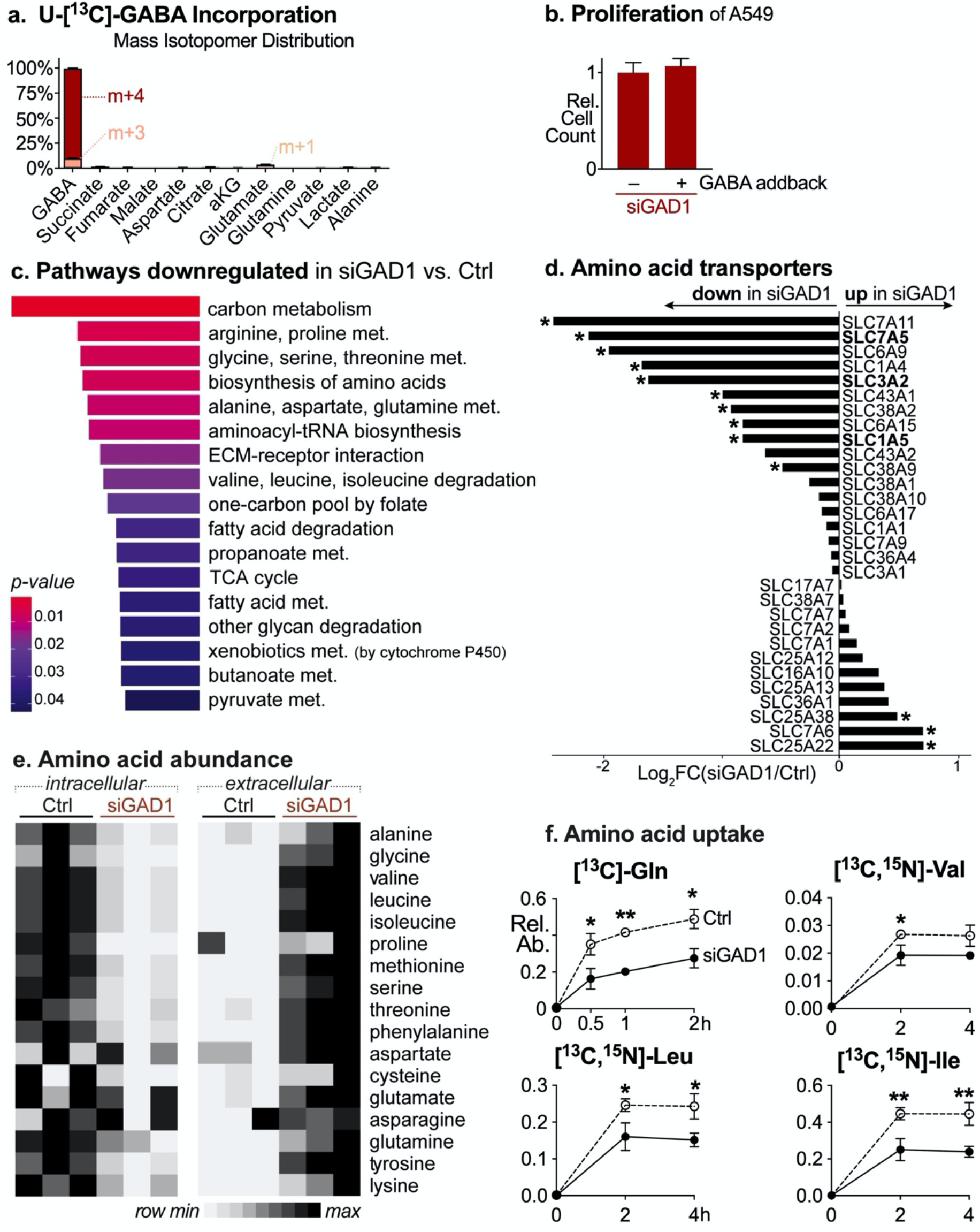
GAD1 regulates amino acid homeostasis independent of GABA metabolism. **A**. Distribution of mass isotopologues in A549 cells incubated with 1 mM U-[^13^C]-GABA for 16 h before extraction. **B**. Proliferation of A549 cells transfected with siRNA targeting GAD1 and treated without (–) or with (+) GABA addback for 96 h. Data are relative to cell counts in GABA-free culture conditions. Data are represented as mean ± SD for replicates (n=4). **C**. Gene set enrichment analysis using RNA-sequencing data for significantly downregulated KEGG pathways in H1299 cells transfected with siRNA targeting GAD1 in comparison to cells transfected with control siRNA (q < 0.05 using the Benjamini-Hochberg method). **D**. Waterfall plot representing the Log2 fold change in amino acid transporter mRNA expression in H1299 cells transfected with either control siRNA (Ctrl) or siRNA targeting GAD1 (siGAD1). *, q < 0.05 using the Benjamini-Hochberg method. **E**. Heat maps respresenting intracellular (*left*) and extracellular (*right*) amino acid abundances in H1299 cells transfected as in D. Values for each metabolite are normalized to a myristic acid internal standard and scaled by row. Data represent three independent cultures. **F**. Kinetics of amino acid uptake in GAD1-deficient cells. H1299 cells were transfected as in D, followed by incubation with either 2 mM U-[^13^C]-Gln, 0.094 g/L U-[^13^C, ^15^N]-valine, 0.105 g/L U-[^13^C, ^15^N]-leucine or 0.105 g/L U-[^13^C, ^15^N]-isoleucine. Cell extracts were harvested at the time points indicated and intracellular levels of the indicated metabolites were determined relative to an internal standard (D-myristic acid).

To explore the impact of GAD1 expression on NSCLC cell proliferation, we performed transcriptional profiling via RNA-sequencing of cells transfected with either control siRNAs or siRNAs targeting GAD1. GAD1 knockdown promoted widespread changes in the transcriptome of A549 cells (17,344 genes detected, 1867 decreased and 1881 increased; q value < 0.05). Gene Set Enrichment Analysis (GSEA) of KEGG downregulated pathways (Figure 2C) highlighted a striking signature of GAD1 knockdown cells characterized by reduced expression of transcripts involved in central carbon metabolism and amino acid homeostasis. Indeed, expression of a third of all amino acid transporters was significantly reduced in cells upon GAD1 silencing (Figure 2D), including the branched chain amino acid (BCAA) transporter LAT1 (*SLC7A5* and *SLC3A2*) and the glutamine transporter ASCT2 (*SLC1A5*). Consistent with this, intracellular levels of 17 amino acids were decreased in GAD1-deficient cells, with corresponding elevation of these amino acids in the extracellular media, suggesting a deficiency in transport (Figure 2E). GAD1 knockdown cells also displayed reduced uptake of both glutamine (U-[^13^C]-Gln) and BCAAs (U-[^13^C,^15^N]-valine, U-[^13^C,^15^N]-leucine, and U-[^13^C,^15^N]-isoleucine) (Figure 2F), confirming that deficiency in GAD1 impairs amino acid import.

To support mRNA translation and unchecked proliferation, tumor cells require a continuous supply of amino acids for biosynthesis (Wortel et al., 2017). Amino acid levels also influence intracellular signalling by regulating activation of the mammalian target of rapamycin complex 1 (mTORC1) (Tokunaga et al., 2004). mTORC1 coordinates cell proliferation and growth by regulating anabolic metabolic pathways, including protein and nucleotide biosynthesis, and is itself regulated by growth factor and nutritional inputs (Ben-Sahra and Manning, 2017). Therefore, we hypothesized that reduced intracellular amino acid levels (in particular impaired leucine transport via LAT1) may impact cellular proliferation in GAD1-deficient cells through reduced mTORC1 activity (Kandasamy et al., 2018). Baseline mTORC1 activity was reduced in GAD1-deficient H1299 cells and further depleted by amino acid starvation, as determined by phosphorylation of the downstream mTORC1 targets S6K (pT389), 4E-BP1 (pT37/46), and ULK1 (pS757), as well as phosphorylation of the S6K target ribosomal S6 protein (pS235/236) (Figure 3A). The rapid loss of mTORC1 activity upon amino acid withdrawal suggests that GAD1-deficient cells remain sensitive to extracellular amino acid levels. Yet, even after serum stimulation, GAD1 deficiency resulted in reduced growth factor-mediated mTORC1 activation, with lower levels of ULK, S6K, and rS6 phosphorylation observed in cells with GAD1 knockdown, particularly at later timepoints (>30 min post-serum stimulation) (Figure 3B).

**Figure 3:**
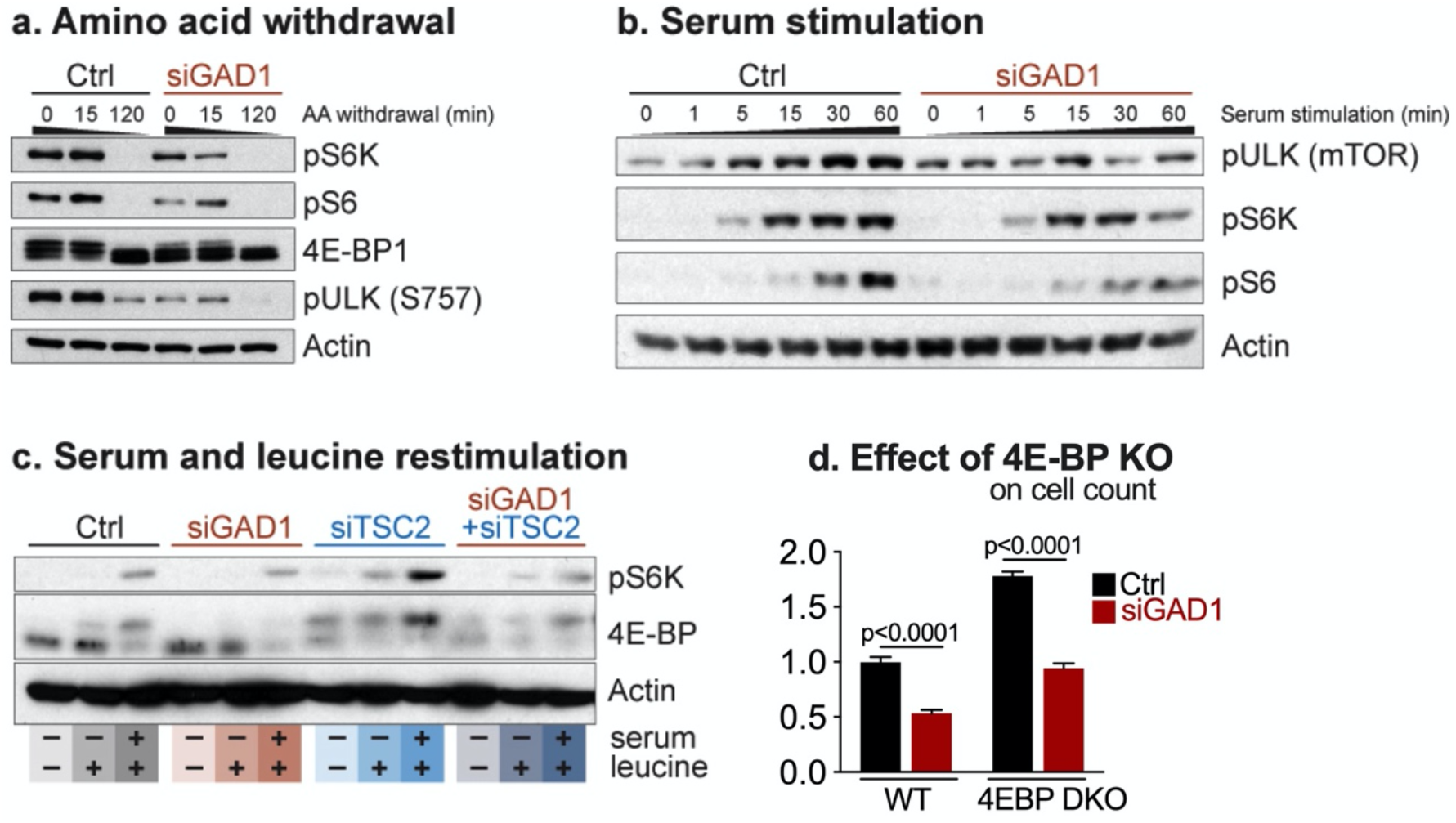
GAD1 regulates mTORC1 activity through control of amino acid supply. **A–C**. mTORC1 activity in GAD1-deficient cells. **A**. H1299 cells were transfected with either control siRNA (Ctrl) or siRNA targeting GAD1 (siGAD1) and mTORC1 activity following amino acid (AA) withdrawal was determined by immunoblot for phospho-p70 S6K (T389), phospho-S6 (S235/236), 4EBP1 and phospho-ULK (S757). Actin was used as a control for protein loading. **B**. H1299 cells transfected as in **A** were serum starved for 50 min, followed by restimulation with serum for the indicated time points. Cell lysates were analysed for ULK, S6K, and rS6 phosphorylation by immunoblot as in **A. C**. HEK293 cells were transfected with either control siRNA (Ctrl) or siRNAs targeting either GAD1 (siGAD1), TSC2 (siTSC2), or both GAD1 and TSC2 (siGAD1+siTSC2). Serum- and BCAA-starved cells were stimulated for 10 minutes with combinations of dialyzed FBS (10% serum) and leucine (400 μM), followed by extraction for immunoblotting. Levels of phosphorylated S6K, total 4E-BP1, and actin are shown. **D**. Effect of GAD1-deficiency on mTORC1-dependent cell proliferation. Wild-type (WT) and 4EBP1/2-deficient (4EBP DKO) MEFs were transfected with either control siRNA (Ctrl) or siRNA targeting GAD1 (siGAD1) and cell counts after 72 h of proliferation were determined by cell counting. Data are presented as mean ± SD for replicates (n=20).

One of the key upstream regulators of mTORC1 activity is the tuberous sclerosis complex (TSC), which activates the GTPase Rheb to hydrolyze the GTP to GDP (Garami et al., 2003). In its GDP-bound state, Rheb cannot activate mTORC1 (Saucedo et al., 2003; Tee et al., 2003). However, locked in its GTP-bound state—caused by deficiency in the TSC component TSC2— Rheb promotes constitutive mTORC1 activity (Garami et al., 2003). As expected, knockdown of TSC2 promoted elevated S6K and 4E-BP phosphorylation in the absence of exogenous signals and increased mTORC1 signalling upon re-addition of serum and, specifically, leucine to culture conditions (Figure 3C) (Sancak et al., 2008; Smith et al., 2005). However, silencing GAD1 reduced mTORC1 signalling in TSC2-deficient cells (Figure 3C). These data suggest that GAD1 acts upstream of TSC2 in regulating mTORC1 activity, likely through regulation of amino acid uptake, and that GAD1 loss is dominant over TSC2-mediated control of mTORC1. To further investigate the relationship between GAD1 and mTORC1-dependent proliferation, we employed 4E-BP1/2 double knockout (DKO) MEFs, which display increased basal proliferative capacity due to their role in translating mRNAs involved in proliferation and cell cycle progression (Dowling et al., 2010). Knockdown of GAD1 ablated the proliferative advantage of 4E-BP DKO MEFs, suggesting that GAD1 is required for proliferation regardless of 4E-BP status (Figure 3D). Together these data indicate a dominant effect of GAD1 loss on the mTORC1 pathway, which likely occurs at the level of amino acid supply rather than by direct interaction with the mTORC1 pathway itself.

Collectively, our data suggest that reduced expression of GAD1 results in intracellular amino acid imbalance. We have shown that one of the major regulatory pathways that controls cellular amino acid levels—the mTORC1 pathway—is negatively impacted by GAD1 loss. We next investigated whether the second major regulatory pathway—the general amino acid control (GAAC) pathway—was also disrupted upon reduced expression of GAD1 (Kandasamy et al., 2018). The major transcription factor controlling the GAAC pathway is ATF4, which is induced by amino acid deprivation. A member of the CREB/ATF family of bZIP transcription factors, ATF4 functions as a master regulator of amino acid uptake and biosynthesis in response to cellular stress (Harding et al., 2003). ATF4 has also been shown to have a pro-tumorigenic role in cancer, promoting xenograft tumor growth and metastasis (Dey et al., 2015; Ye et al., 2014), and has been shown to upregulate genes correlated with poor prognosis in NSCLC (DeNicola et al., 2015). More recently, ATF4 has been shown to maintain the homeostatic balance between mRNA translation supply and demand under non-stressed conditions by tuning the expression of the many enzymes and transporters required for delivering amino acids to the translation machinery (Park et al., 2017). Given that GAD1-deficient cells have a plentiful supply of amino acids (but fail to import them), we investigated the relationship between GAD1 loss and ATF4.

Analysis of genes downregulated upon GAD1 knockdown revealed significant overlap with genes downregulated in NSCLC cells transfected with siRNA targeting the transcription factor ATF4 (Igarashi et al., 2007) (Figure 4A). This signature (18 common genes) included several metabolic enzymes and transporters involved in amino acid metabolism including the BCAA transporter *SLC7A5* and *SLC3A2* (LAT1) and the serine biosynthesis pathway enzymes PHGDH and PSAT1 (validated in Figures 4B–C and S3A). Silencing either GAD1 or ATF4 reduced *SLC7A5* mRNA expression (Figure 4B) and LAT1 protein expression (Figure 4C) in NSCLC cells. Acute knockdown of ATF4 also led to a comparable proliferative defect as GAD1 knockdown (Figure 4D). Upon amino acid stress, the eukaryotic initiation factor eIF2α is phosphorylated by GCN2 (general control non-derepressible 2) leading to suppression of global protein synthesis but an enhancement of ATF4 protein levels (Wortel et al., 2017). Levels of phospho-GCN2 and phospho-eIF2α were enhanced upon amino acid withdrawal and suppressed by the presence of extracellular glutamine in both control and GAD1 knockdown cells (Figure S3B). eIF2α phosphorylation levels were, if anything, slightly elevated in GAD1 knockdown cells upon amino acid withdrawal, suggesting amino acid sensing was intact and that GAD1 impacts engagement of the downstream ATF4-dependent transcriptional program that normally regulates amino acid homeostasis under basal growth conditions.

**Figure 4:**
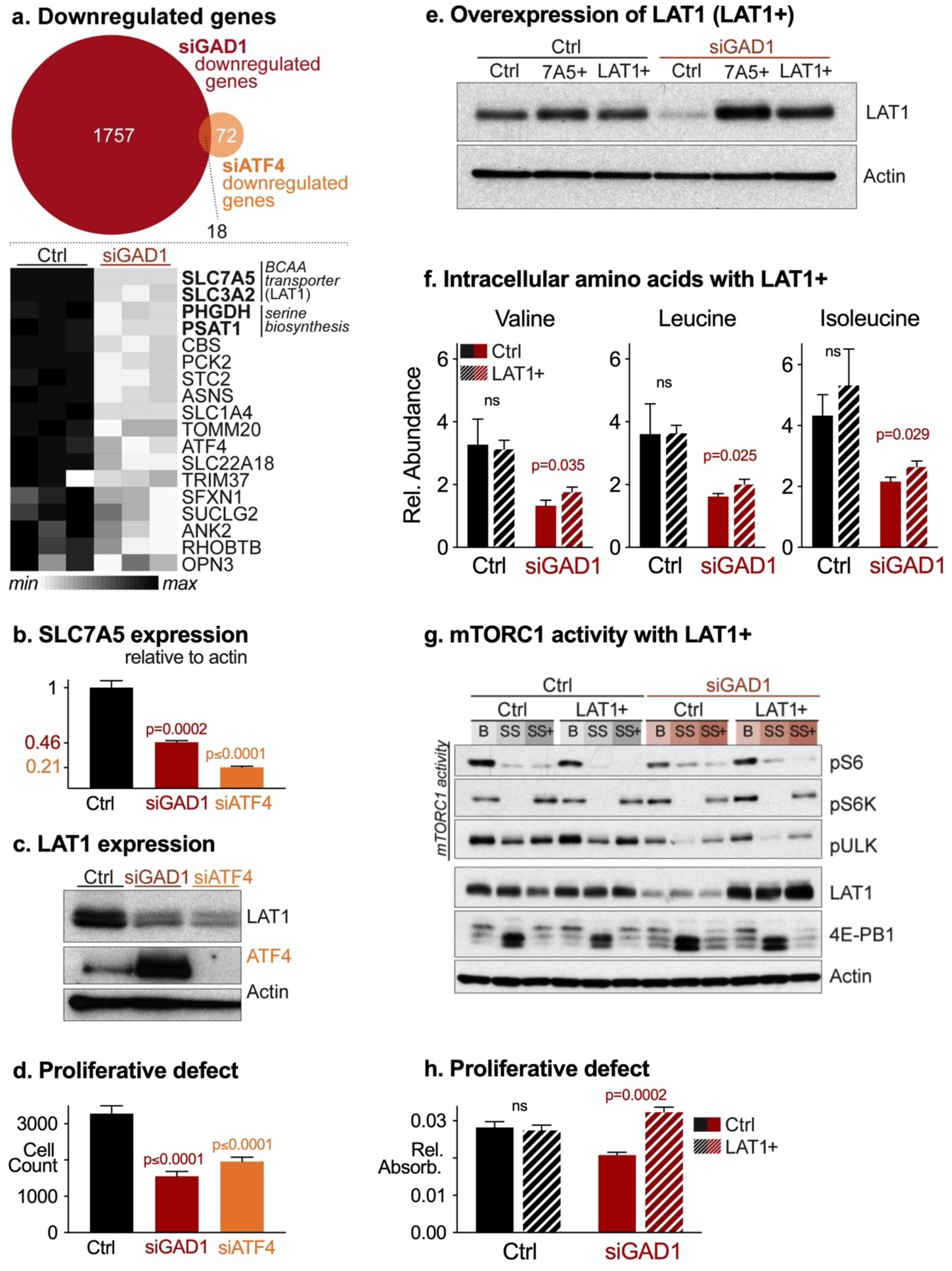
GAD1 regulates ATF4-dependent amino acid homeostasis. **A**. Top, Venn diagram depicting downregulated genes (1757 genes) from H1299 cells transfected with siRNA targeting GAD1 (siGAD1), relative to cells transfected with control siRNA (Ctrl), as determined by RNA sequencing. Overlap of GAD1-regulated genes with the Igarashi ATF4 target gene signature is shown. Bottom, heat map analysis of the 18 ATF4-dependent genes downregulated in GAD1-deficient H1299 cells. **B**. Relative expression of *SLC7A5* mRNA in H1299 cells transfected with either control siRNA (Ctrl) or siRNAs targeting GAD1 (siGAD1) or ATF4 (siATF4). Transcript levels were determined relative to ACTB mRNA levels. Data are represented as mean ± SD (n=3). **C**. Immunoblot of LAT1 and ATF4 expression in H1299 cells transfected as in B. Actin was used as a control for protein loading. **D**. Proliferation of GAD1- and ATF4-deficient H1299 cells. H1299 cells were transfected as in B, and cell number after 48 h of proliferation was determined by cell counting. Data are represented as mean ± SD for replicates n=60. **E**. Immunoblot for LAT1 expression in H1299 cells with ectopic expression of either *SLC7A5* alone (7A5+) or *SLC7A5* and *SLC3A2* (LAT1+). Cells were transfected with control siRNA (Ctrl) or GAD1 siRNAs (siGAD1) followed by expression of 7A5+ or LAT1+ for 48h prior to extraction of protein lysates for immunoblotting. Actin was used as a control for protein loading. **F**. Abundance of branched chain amino acids (valine, leucine, isoleucine) in H1299 cells expressing LAT1. Control (Ctrl) or *SLC7A5/SLC3A2*-expressing (LAT1+) H1299 cells were transfected with control siRNA (Ctrl) or GAD1 siRNA (siGAD1), and intracellular amino acid levels were determined 48 h later. Metabolite abundance is presented relative to an internal standard (D-myristic acid). Data are represented as mean ± SD from triplicate plates (n=3 for each group). **G**. Immunoblot for mTORC1 activity (phospho-p70 S6K (T389), phospho-S6 (S235/236), 4E-BP1 (pT37/46), phospho-ULK (S757)) and LAT1 expression in H1299 cells transfected as in **F**. Cells were extracted either under basal growth conditions (B), serum starvation (SS, 50 min), or serum addback (SS+, serum starvation followed by 10 min serum stimulation). **H**. Proliferation of GAD1-deficient cells expressing LAT1. H1299 cells transfected as in F were plated and relative absorbance measured (as proxy for cell count) after 48h of proliferation. Data are represented as mean ± SD for replicates n=15

Lastly, we considered whether the observed defects in the control of amino acid homeostasis were responsible for the proliferative defect of GAD1-deficient cells (Figure 1D). To address this directly, we ectopically expressed the BCAA transporter *SLC7A5*/*SLC3A2* (LAT1) in GAD1-deficient cells to restore BCAA transport (Figure 4E) and assessed the impact on cellular metabolic and proliferative capacity. Overexpression of LAT1 in GAD1 knockdown cells led to a small but significant increase in intracellular BCAA levels (Figure 4F) and restored S6K and rS6 phosphorylation under basal conditions (Figure 4G). Finally, we found that expression of LAT1 was sufficient to enhance proliferation of GAD1-deficient cells (Figure 4H). These data suggest that reduced amino acid supply, downstream of defective ATF4 signalling is responsible for the proliferative defect observed in cells lacking GAD1.

Several recent papers have explored the regulation of GAD1 expression in tumors. DNA methylation of the GAD1 promoter has been shown to promote GAD1 activation in cancer cells (Yan et al., 2016) and epigenetic upregulation of GAD1 has been shown in the brain metastatic environment (Schnepp et al., 2017). GAD1 methylation positively correlated with mRNA expression in lung adenocarcinoma tissue samples and GAD1 mRNA and protein expression were also significant prognostic indicators (Tsuboi et al., 2019). Beyond lung cancer, GAD1 may play a role in metastasis in glioma (Schnepp et al., 2017) and oral squamous cell carcinoma (Kimura et al., 2013). Our data indicate that GAD1 expression is elevated in a number of tumor types (Figure 1B) and high GAD1 expression correlates with worse outcomes in lung cancer (Figure 1C). Although there is evidence that tumors of neuronal origin might engage the GABA shunt (Ippolito and Piwnica-Worms, 2014; Schnepp et al., 2017), to date there has been little mechanistic evidence defining a role for GABA shunt enzymes, such as GAD1, in tumor development.

Our results reveal a novel regulatory pathway mediated by the enzyme GAD1 that influences cancer cell proliferation through control of amino acid homeostasis. While GAD1, as part of the GABA shunt metabolic pathway, may provide an alternative method of glutamine anaplerosis in plants (Shelp et al., 2012) and neuronal tissues (Hertz, 2013), our data suggest it clearly occupies an alternative role in NSCLC cells in maintaining cellular amino acid homeostasis. We show here that silencing GAD1 leads to a global reduction in amino acid transporters, including the BCAA transporter LAT1, resulting in reduced amino acid import and mTORC1 activation. Non-canonical functions have been demonstrated for other metabolic enzymes. For example, GAPDH and PKM2 have been shown to perform multiple functions beyond their established roles in glycolysis (Chang et al., 2013; Gupta and Bamezai, 2010). To our knowledge, only Kimura et al. have proposed a role for GAD1 independent of the GABA shunt metabolic pathway, proposing that GAD1 promotes cellular invasiveness and migration by controlling β-catenin localization and subsequent Wnt signal transduction (Kimura et al., 2013). While our data cannot rule out contributions of Wnt signalling to the metabolic alterations induced by GAD1 loss, our data prominently indicate that GAD1 has a non-canonical function: maintaining cellular growth and proliferation programs through the regulation of amino acid homeostasis.

Our data suggest that GAD1 is required for maintenance of basal amino acid homeostasis through a mechanism involving the stress-responsive transcription factor ATF4. While it plays a key role in regulating stress responses to amino acid withdrawal, ATF4 has also been shown to regulate amino acid homeostasis under non-stressed conditions (Park et al., 2017; Quirós et al., 2017). Park *et al*. report that basal levels of ATF4 appear to target different genes than those induced by endoplasmic reticulum (ER) stress when levels of ATF4 are significantly elevated (Park et al., 2017). Similar to our results with GAD1 knockdown (Figure 2E and F), they show that in basal conditions, loss of ATF4 reduced the uptake of 15 amino acids, demonstrating that lost ATF4 activity is sufficient to reduce basal amino acid uptake (Park et al., 2017). They also note that amino acid uptake remained sensitive to mTOR activity in ATF4-null cells, indicating that mTOR controls amino acid uptake through both ATF4-dependent and -independent mechanisms. We propose that GAD1 loss interferes with this program, preventing mTORC1 activation and subsequent ATF4 mediated transcription of amino acid transporters and metabolic enzymes involved in amino acid metabolism. In support of this, we observe 4E-BP hypophosphorylation in GAD1 knockdown cells at baseline, the extent of which is similar to control cells following a period of amino acid withdrawal (Figure 3A). This further supports our hypothesis that GAD1 deficiency induces a state of amino acid starvation that impacts cell proliferation. Disruption of the interplay between mTORC1 and ATF4 could be particularly pertinent in NSCLC, given that oncogenic KRAS (mutated in ∼30% of NSCLC) has been shown to regulate amino acid homeostasis and cellular response to nutrient stress via ATF4 (Gwinn et al., 2018). Future work will focus on the mechanism(s) by which GAD1 regulates ATF4 function.

In summary, our work has revealed a novel regulatory mechanism for amino acid homeostasis in tumor cells mediated by the GABA shunt enzyme GAD1. GAD1 is overexpressed in several cancers, and our results suggest that this enzyme exerts distinct functions from its role in GABA metabolism that impact cancer cell metabolism and proliferation.

## Acknowledgments

We thank D. Avizonis, G. Bridon, and L. Choinière from the McGill University Metabolomics Core Facility for technical assistance. R.G.J. is supported by grants from the Canadian Institutes of Health Research (CIHR, MOP-93799) and the Terry Fox Research Institute (TFRI, TFRI-239585). E.E.V is supported by Cancer Research UK (C18281/A19169) and by a Diabetes UK RD Lawrence Fellowship (17/0005587).

## Methods

### Cell lines and cell culture

H1299 and A549 cells were obtained from the ATCC (Manassas, VA, USA). MEF cells deficient for 4EBP1 and 4EBP2 and TSC2 were kindly provided by Masahiro Morita and Nahum Sonenberg (Dowling et al., 2010; Morita et al., 2013). Cells were cultured in growth medium (DMEM [A549, HEK293 and MEF cells] or RPMI medium [H1299 cells]) supplemented with 10% fetal bovine serum (FBS), penicillin, streptomycin, glutamine, and non-essential amino acids (for H1299 cells). Cells were grown at 37°C in a humidified atmosphere supplemented with 5% (v/v) CO_2_. Amino acid withdrawal was stimulated by replacing culture medium with Eagle’s Buffered Salt Solution (EBSS) containing 25 mM glucose for indicated times. For amino acid stimulation experiments, cells were starved of serum and branched chain amino acids for 50 min, followed by restimulation with combinations of dialyzed FBS (10% serum) and leucine (400 μM) for 10 min before extraction. Transient knockdown of GAD1, ATF4, and TSC2 was achieved using SMARTpool ON-TARGETplus siRNA reagent (composed of four individual siRNAs for each target) from GE Dharmacon. Briefly, 50 nM siRNA was incubated in a cell culture plate for 20 min with RNAimax in OptiMEM medium (Life Technologies) onto which the cells were seeded. Cells were transfected a second time 48 h later and plated for assays the following day. Unless otherwise stated cells were harvested for RNA-sequencing, metabolomics and western blotting 48 h after seeding. Overexpression of LAT1 was achieved by transfecting cells with constructs pcDNA3-*SLC7A5* and pcDNA3-*SLC3A2* using Lipofectamine 2000 as per the manufacturer’s recommendations.

### Cell proliferation and cell counting assays

Cells were either seeded in 96-(8000 cells/well) or 384-well plates (500 cells/well) in growth medium. At time points indicated in the figure legends, cells were fixed with 4% formaldehyde. Cells were either stained with crystal violet as described previously (Vincent et al., 2014) or stained with Hoechst DNA stain, and cell number was determined by nuclei counting. Images were taken using an Operetta High Content Imaging System and analyzed using Harmony High Content Imaging and Analysis Software, both from Perkin Elmer (Waltham, MA, USA).

### Immunoblotting and quantitative real-time PCR

Cells were lysed in modified CHAPS buffer (10 mM Tris·HCl, 1 mM MgCl2, 1 mM EGTA, 0.5 mM CHAPS, 10% glycerol, 5 mM NaF) or AMPK lysis buffer (Faubert et al., 2013) supplemented with the following protease additives: protease and phosphatase tablets (Roche), DTT (1 μg/mL), and benzamidine (1 μg/mL). Cleared lysates were resolved by SDS-PAGE, transferred to nitrocellulose, and incubated with primary antibodies. Primary antibodies to S6 ribosomal protein (pS235/236), 4E-BP1 (pT37/46), p70 ribosomal S6 Kinase (pT389), ULK (pS757), GCN2 (phospho-Thr898 and total), eIF2α (phospho-S51) LAT1, ATF4, and actin, as well as HRP-conjugated anti-rabbit secondary antibodies were obtained from Cell Signaling Technology (Danvers, MA, USA). qPCR was performed as described previously (Faubert et al., 2013). Primer sequences are listed in Table S1.

### Metabolite profiling by GC-MS and LC-MS

Cellular metabolites were extracted and analyzed by either gas chromatography-mass spectrometry (GC-MS) or liquid chromatography (LC)-MS using protocols described previously (McGuirk et al., 2013; Vincent et al., 2015). Briefly, for GC-MS, metabolite extracts were derived using N-(tert-butyldimethylsilyl)-N-methyltrifluoroacetamide (MTBSTFA). D-myristic acid (750 ng/sample) was added as an internal standard to metabolite extracts, and metabolite abundances are expressed relative to the internal standard and normalized to cell number. LC was performed using a 1290 Infinity ultraperformance LC system (Agilent Technologies) equipped with a Scherzo 3-μm, 3.0 × 150 mm SM-C18 column (Imtakt). LC-MS analysis was performed on an Agilent 6540 ultra-high definition (UHD) Accurate-Mass Q-TOF mass spectrometer (Agilent). For stable isotope tracer analysis (SITA) experiments, cells were cultured with U-[^13^C]-glutamine, U-[^13^C]-GABA (Cambridge Isotopes), [^13^C, ^15^N]-valine, [^13^C, ^15^N]-leucine, or [^13^C, ^15^N]-isoleucine (Sigma Aldrich) for the indicated times. Mass isotopomer distribution was determined using a custom algorithm developed at McGill University (McGuirk et al., 2013).

### RNA Sequencing

RNA was extracted from H1299 cells following transfection with siRNA. cDNA synthesis and library construction were conducted as described previously (Jha et al., 2015). Libraries were sequenced at the Centre for Applied Genomics (SickKids, Toronto) using a HiSeq 2500 (Illumina) using 50 × 25 bp pair-end sequencing. GSEA on RNA-seq data was conducted using the gage function and non-parametric Kolmogorov-Smirnov test from the GAGE R Bioconductor package (Luo et al., 2009). The Igarashi *et al*. ATF4 target gene signature MSigDB has been previously described (Igarashi et al., 2007).

### TCGA Data Analysis

TCGA data from 14 tumor studies that contained mRNA expression for at least five normal and five tumor samples were included in the analysis. Data were accessed using the R-interface cgdsr to the cBio Cancer Genomics Portal (Cerami et al., 2012; Gao et al., 2013). For each dataset, *GAD1, ABAT*, and *ALDH5A* expression levels between normal and tumor tissue samples were compared.

### Survival analysis

The Kaplan–Meier plotter (http://kmplot.com) (Győrffy et al., 2013) was used to assess the effect of *GAD1, ABAT* and *ALDH5A* expression levels on overall lung cancer survival. Lung cancer patients were divided into high and low expression of *GAD1, ABAT*, and *ALDH5A* by median value of mRNA expression.

### Statistical Analysis

Data are presented as mean ± SD for technical replicates, or mean ± SEM for biological replicates, and analyzed using unpaired Student’s *t* test. Statistical significance is indicated in all figures by the following annotations: *, *p* < 0.05; **, *p* < 0.01; ***, *p* < 0.001, unless otherwise noted.

## Supplemental Information

**Supplementary Figure 1:**
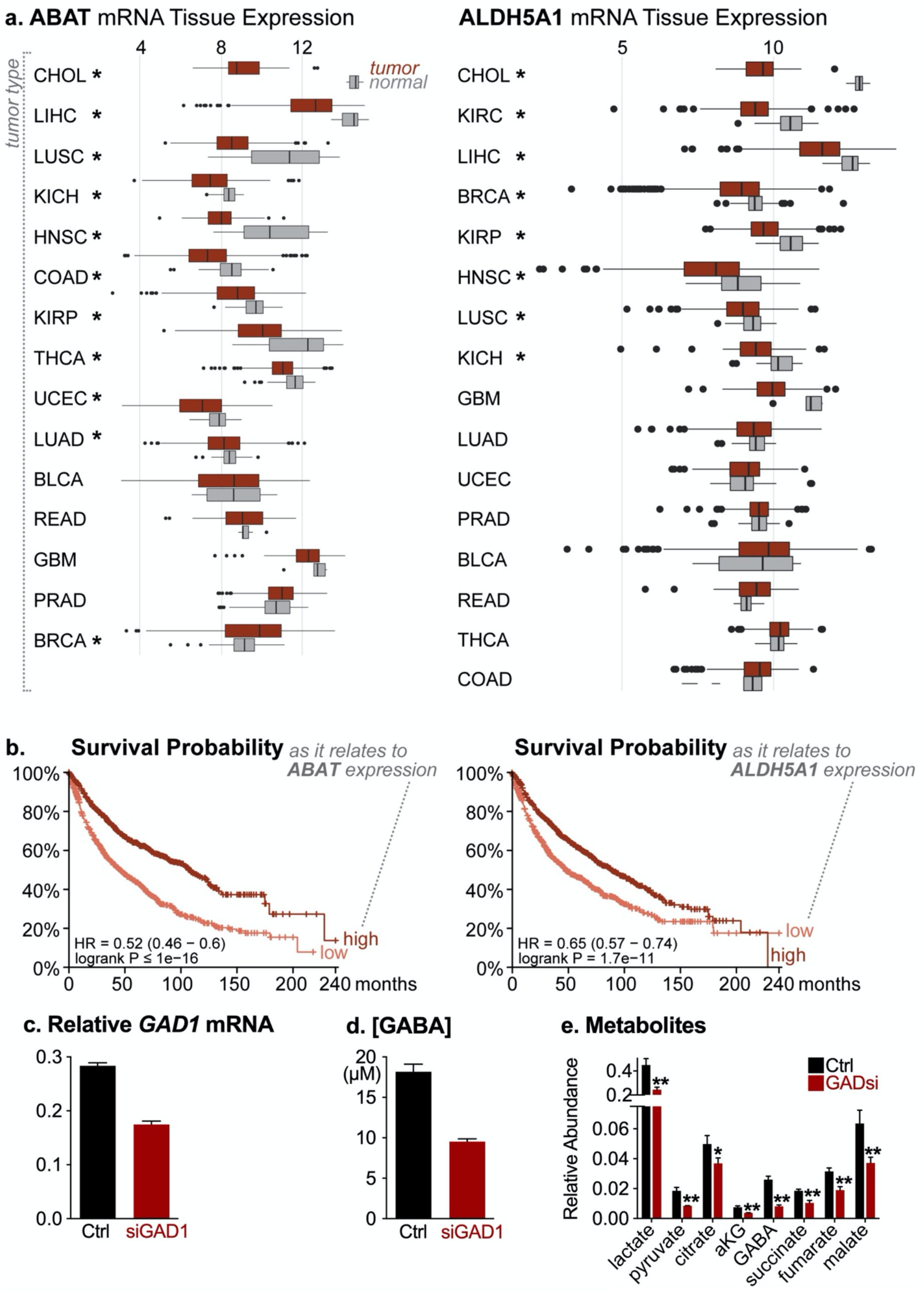
**A**. Boxplot of data taken from The Cancer Genome Atlas (TCGA) database representing tumour tissue and normal tissue mRNA expression of *ABAT* and *ALDH5A*. Data presented as Log2 expression. Data set sorted by t-test statistical value, with large down-regulation in tumours at the top (* denotes adjusted p < 0.01). CHOL: Cholangiocarcinoma, LICH: XXX, LUSC: Lung Squamous Cell Carcinoma, KICH: Kidney Chromophobe, HNSC: Head-Neck Squamous Cell Carcinoma, COAD: Colon Adenocarcinoma, KIRP: Cervical Kidney Renal Papillary Cell cCarcinoma, THCA: Thyroid Cancer, UCEC: Uterine Corpus Endometrial Carcinoma, LUAD: Lung Adenocarcinoma, BLCA: Urothelial Bladder Carcinoma, READ: Rectum Adenocarcinoma, GBM: Glioblastoma, PRAD: Prostate Adenocarcinoma, BRCA: Breast Invasive Carcinoma, KIRC: Kidney Renal Clear Cell Carcinoma. **B**. Kaplan–Meier overall survival curve in lung cancer patients stratified into high (n=963) or low (n=963) expression of *ABAT* and stratified into high (n=962) or low (n=964) expression of *ALDH5A1* mRNA using the median expression value as the cut-off point. **C**. Relative expression of *GAD1* mRNA in H1299 cells as determined by qPCR. H1299 cells were transfected with either control (Ctrl) or siRNA targeting GAD1 (siGAD1) and cultured for 72 h before mRNA extraction. Transcript levels were determined relative to *ACTB* mRNA levels. Data are represented as mean ± SD for triplicate samples (n=3). **D**. Adundance of GABA in H1299 cells transfected as in **C**. Data are represented as mean ± SEM of three independent cultures. **E**. Intracellular metabolite levels in H1299 cells transfected with either control siRNA (siCtrl) or siRNA targeting GAD1 (siGAD1). Values for each metabolite abundance are relative to a myristic acid internal standard. These data are also displayed in the heat map in Figure 1E.

**Supplementary Figure 2:**
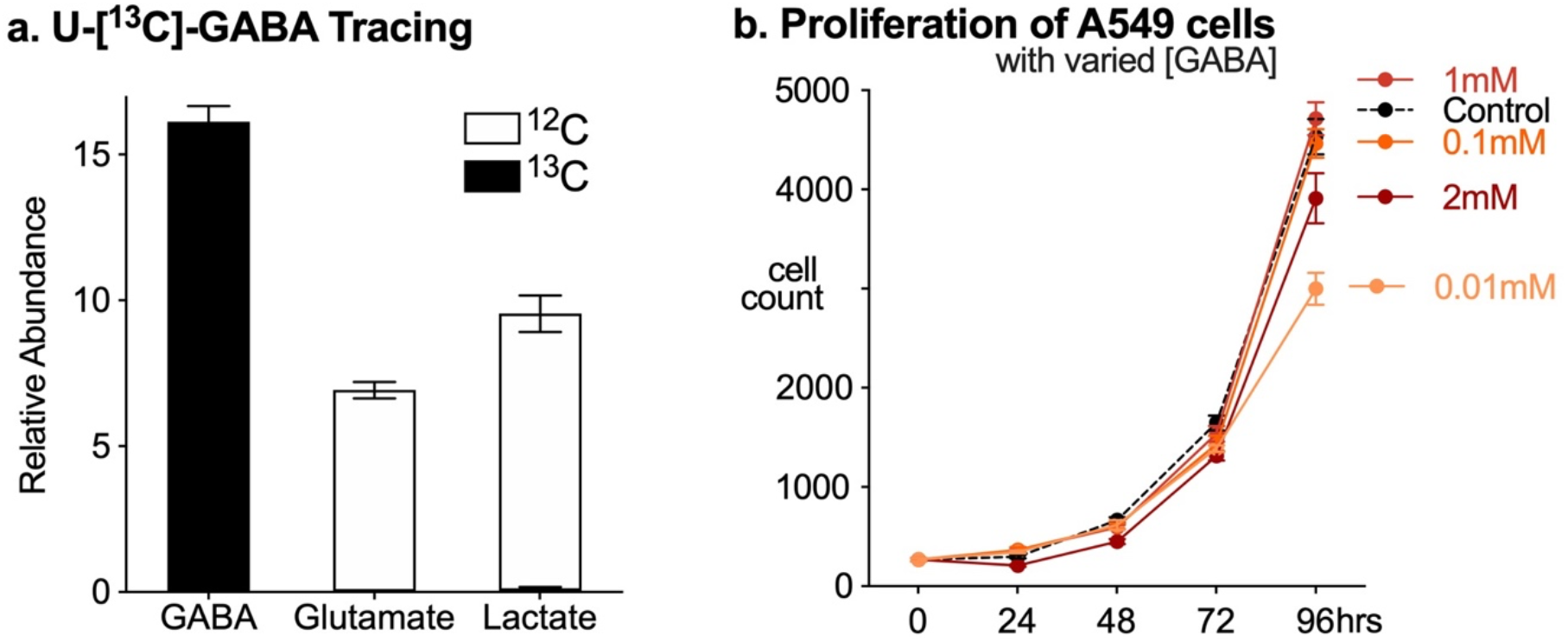
**A**. Relative abundance of GABA, glutamate, and lactate in A549 cells incubated with 1 mM U-[^13^C]-GABA for 16 h before extraction. Abundance of ^13^C-GABA-derived metabolites are shown in black, while unlabeled (^12^C) metabolite abundances are shown in white. Metabolite abundance data are presented relative to an internal standard (D-myristic acid). Data presented as mean ± SEM of three independent cultures (n=3). **B**. Proliferation of A549 cells cultured with varying concentrations of GABA over time. Data are relative to cell counts at the time of plating (0 h). Data are represented as mean ± SD for replicates (n=10 per group).

**Supplementary Figure 3:**
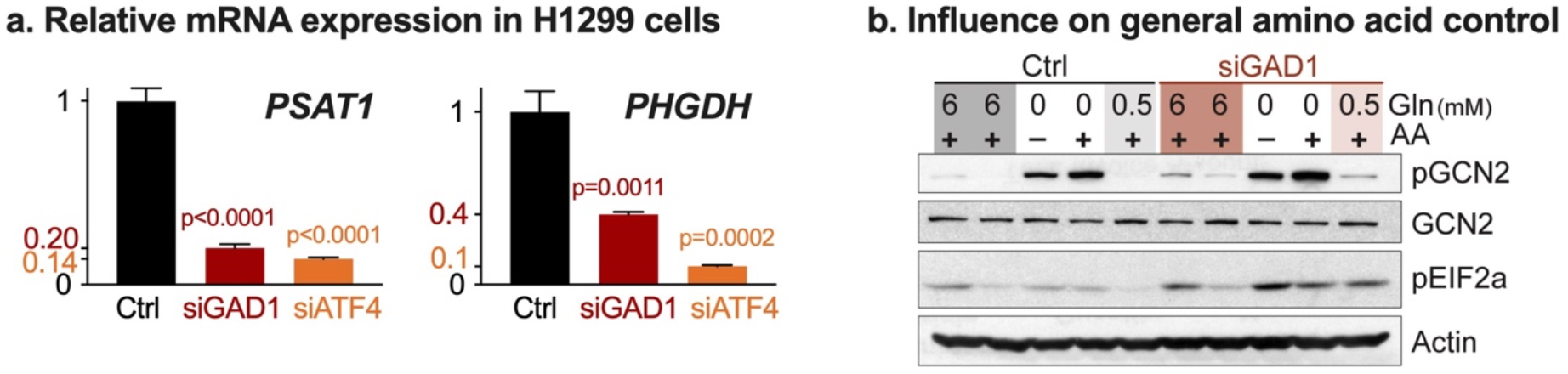
**A**. *PSAT* and *PHGDH* expression in H1299 cells. Relative expression of *PSAT1* and *PHGDH* mRNA in H1299 cells transfected with either control siRNA (Ctrl) or siRNA targeting *GAD1* (siGAD1) or *ATF4* (siATF4). Transcript levels were determined relative to *ACTB* mRNA levels. Data are represented as mean ± SD for replicates (n=3). **B**. Immunoblot for phospho-GCN2 (Thr898), total GCN2, and phospho-eIF2α (Ser51) in H1299 cells. Cells were treated for 4 hours with DMEM media containing 10% FBS and either 0, 0.5 mM, or 6 mM glutamine or amino acid-free medium (-) containing dialyzed 10% FBS. Actin was used as a control for protein loading.

**Table S1:**
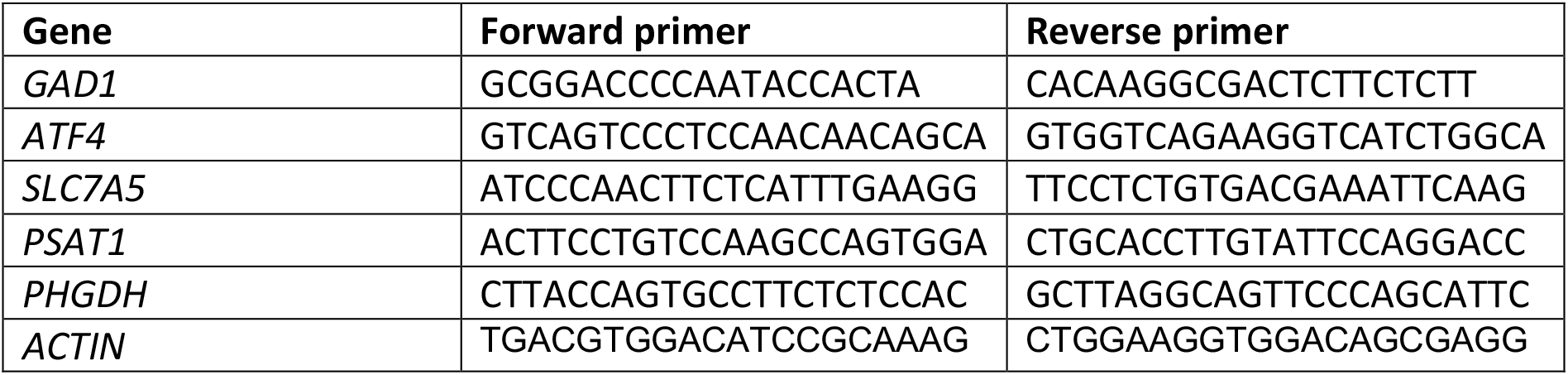
List of qPCR primers

